# Evaluation of new polymer-iodine complexes for the fabrication of medical devices

**DOI:** 10.1101/2021.11.24.469532

**Authors:** Marina López-Álvarez, Herb Ulmer, Nico Klay, Jan Maarten van Dijl

**Affiliations:** Department of Medical Microbiology, University of Groningen, University Medical Center Groningen, Groningen, The Netherlands; X-Infex B.V., The Netherlands

**Keywords:** Povidone-iodine, iodine, polyamide, polyurethane, bacteria, yeast, fungi

## Abstract

Povidone-iodine has been a true success story in the fight against infections by harnessing the antimicrobial and antiviral properties of elemental iodine for water-based applications. However, to date there has been little success in implementing iodine attributes in water-insoluble engineering plastics. Here, we describe the first development of biocidal active polyamide- and polyurethane-iodine complexes at laboratory and commercially relevant scales. These polymer-iodine materials are active against a broad range of microorganisms, including bacteria, yeast and fungi, and can be used as base materials for medical devices. The use of new polymer-iodine complexes for infection prevention in medical devices, such as sutures, catheters and drains, or wound care is expected to have significant positive effects at reducing healthcare-acquired infections. In addition, the materials are expected to find significant applications in other fields, such as air handling with the production of biocidal face masks and air filters to control spread of pathogens.

## 1. Introduction

Healthcare-acquired infections (HAI’s) continue to be a source of countless unneeded deaths and excessive hospital costs. The main sources of HAI’s are pathogenic microorganisms that have entered the body due to contamination and infection, or due to imbalances in the normal microbiota (i.e. dysbiosis) as a consequence of specific treatment conditions or weakened immune defences. Often the HAI forms at the site of a medical device (e.g. a central line catheter) that causes introduction of unwanted microorganisms at a specific body site or becomes the scaffold for unwanted adhesion of bacteria already carried by the patient. In fact, medical devices are a significant source of HAI’s and represent a serious concern in many clinical settings ^[1-3]^. Microbial infections caused by the use of medical devices, such as sutures, catheters, central lines and prostheses result in secondary infections that are often difficult to eradicate and compromise the well-being of the patient. Medical devices provide a perfect nidus for microbial adherence and subsequent formation of biofilms, which are very difficult to eradicate. For these reasons, there is an urgent need to develop and evaluate new materials with biocidal properties in order to reduce the risk of medical device contamination and microbial attachment to said medical devices.

Severe infections may result in removal of medical devices, tissue debridement or, at worse, patient death. Such HAI’s not only worsen patient health, but also increase healthcare costs significantly. In particular, catheters are the most common implants worldwide, with 5 million central venous catheters (CVC) and 30 million urinary catheters being implanted per year, in the USA alone. Both catheter types have been identified as two of the leading origins of HAI’s. The Centers for Disease Control (CDC) reported that approximately 250,000 CVC-related bloodstream infections (BSI) occur each year in the USA at an additional hospitalization cost of ca. $34,500-$56,000 per each infection. Not only is there a huge economic cost, but the CDC estimates that 4% of patients in the USA will contract a HAI during hospitalization, which will result in 1.7 million infections and 99,000 associated deaths.

Further compromising the treatment of HAI’s is the development of “superbugs”, bacteria that have developed resistance to common antibiotics and medications. Not only are healthcare institutions the major source of such “superbugs”, but the people they treat are very young, immunocompromised, elderly and sick, making them the most vulnerable group. The World Health Organization (WHO) has identified antimicrobial resistance as one of the global health threats of the 21^st^ century with estimated casualties amounting to 10 million people/year at a cost of $100 trillion by 2050 if no developments on controlling this health crisis are found.

The water-soluble iodophor complex, povidone-iodine (PVP-I), is a potent broad-spectrum antimicrobial active with excellent antiseptic properties against bacteria, including methicillin-resistant *Staphylococcus aureus* (MRSA) and mycobacteria ^[4]^, fungi ^[5]^, protozoa ^[6]^ and both enveloped and non-enveloped viruses, such as the poliovirus, immunodeficiency viruses ^[6, 7]^, influenza viruses ^[8, 9]^ and coronaviruses: SARS ^[10]^, MERS ^[11]^ and SARS-CoV-2 ^[12]^. Topical formulations of PVP-I have been widely used over the years for disinfection, wound care or treatment of burns ^[8]^ and, despite its common use for more than 60 years, no microbial resistance to PVP-I has ever been reported ^[5]^. However, the PVP-I complex is water soluble and thus its applications in healthcare are generally limited to surface and skin disinfection to minimize the risk of infections.

The research question we addressed in the present study was whether it would be possible to develop water-insoluble iodine-polymer complexes that have similar antimicrobial properties as PVP-I, are relatively simple to manufacture, and can be used for actual medical device fabrication to reduce the risk of infections. More importantly, such new water-insoluble iodine-polymer complexes with biocidal activity could play an important role in reducing the risks of HAI’s in medical devices, equipment and articles by reducing the risk of undesirable microbial contamination and inhibition of microbial adhesion leading to biofilm formation. Therefore, iodine complexes of two polymer families, polyamides (PA) and polyurethanes (PU), were manufactured and the biocidal properties of tubes and foams of PA- and PU-iodine complexes were tested against some of the most commonly found and clinically relevant microorganisms. In brief, we show that the described polymer-iodine complexes can be simply produced by utilizing existing equipment and processes already used to make the corresponding non-iodine polymer materials, and that they have strong bactericidal and fungicidal activity.

## 2. Results

### 2.1. Production of polymer-iodine complexes

Polyamide-Iodine (PA-I) thermoplastic complexes were produced by direct extrusion of the base PA polymer with a suitable iodine source in a twin-screw extruder. The extrusion conditions were dependent on the base polyamide used and the ease that the iodine source is dissolved into the molten PA matrix to generate the PA-I complex. General extrusion temperatures were in the 150-240 °C range, resulting in PA-I complexes that were homogeneous and stable.

Polyurethane-Iodine (PU-I) thermoplastic complexes were produced by direct extrusion of the base PU with a suitable iodine source in a twin extruder. Extruding temperatures of <200 °C were generally effective to generate homogeneous and stable PU-I thermoplastics. PU-I thermosets (cross-linked polymeric matrixes) were produced by first dissolving the suitable iodine source in one of the raw materials for PU fabrication, primarily the polyol or diol raw material. The PU reaction was then conducted as usual to produce the PU-I thermoset. Suitable iodine sources used for the complexation reactions included elemental iodine (I) and povidone-iodine (PVP-I).

The resultant extruded thermoplastic PA- and PU-I tubes were directly tested for antimicrobial activity. The PU-I thermosets (foams) were allowed to cure and then cut into the desired shape for subsequent antimicrobial testing.

### 2.2 Inhibitory effects of thermoplastic PA- and PU-I (P-I) tubes on the growth of various microorganisms

To evaluate the potential growth inhibitory effects of P-I thermoplastic complexes on microorganisms, tubes composed of PA or PU extruded with PVP, PVP-I and/or iodine at various iodine concentrations (**Table 1**) were tested. Eight of the tested tubes (i.e., numbers 3, 4, 6, 14, 15, 16, 17 and 19) showed large clearing inhibition zones for *S. aureus* (**Figure 1**). Tubes 1, 2, 11 and 18 containing lower iodine concentrations showed more localized biocidal activity, with the clearing zones mainly apparent at the point of direct tube contact.

**Table 1.**
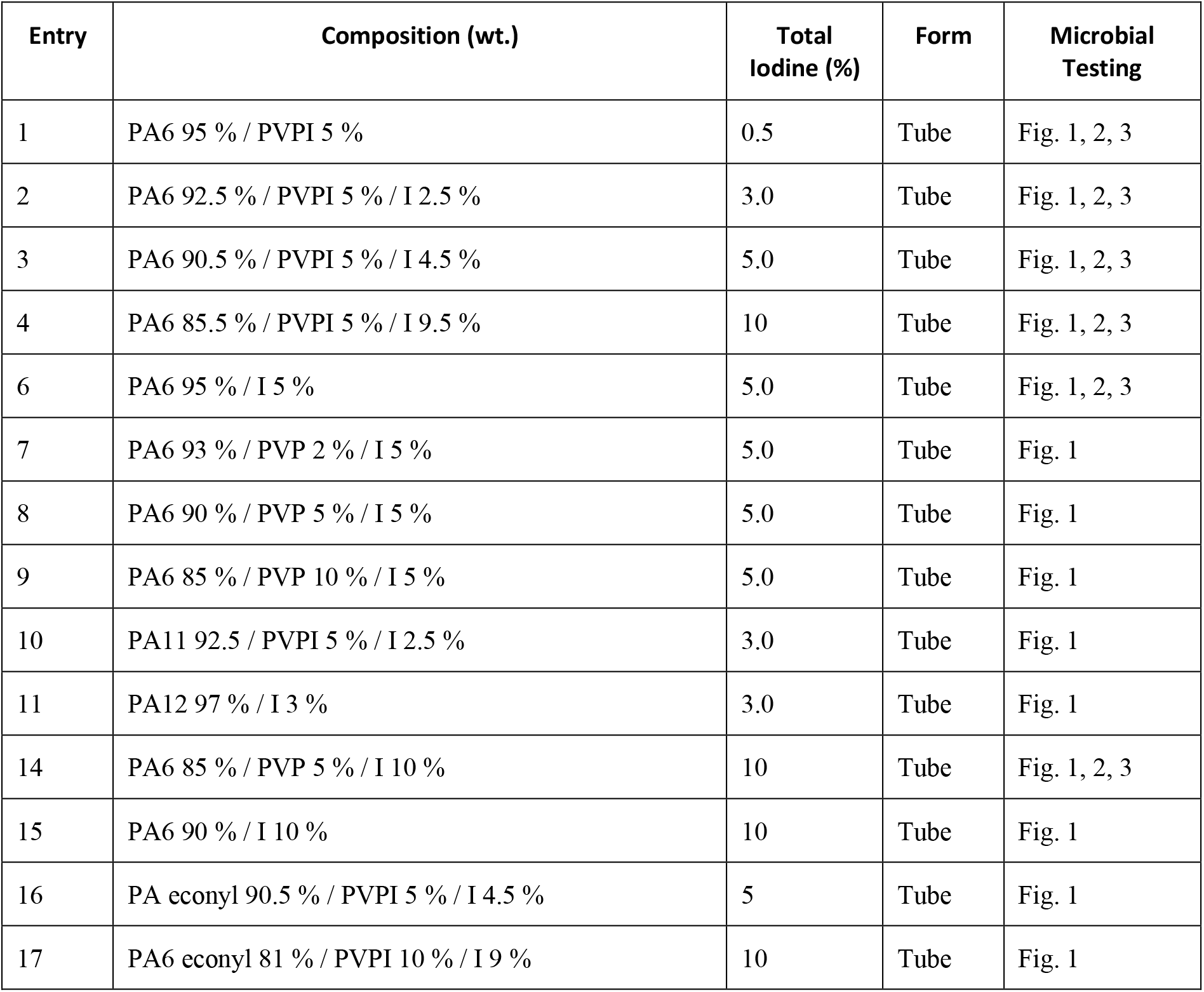
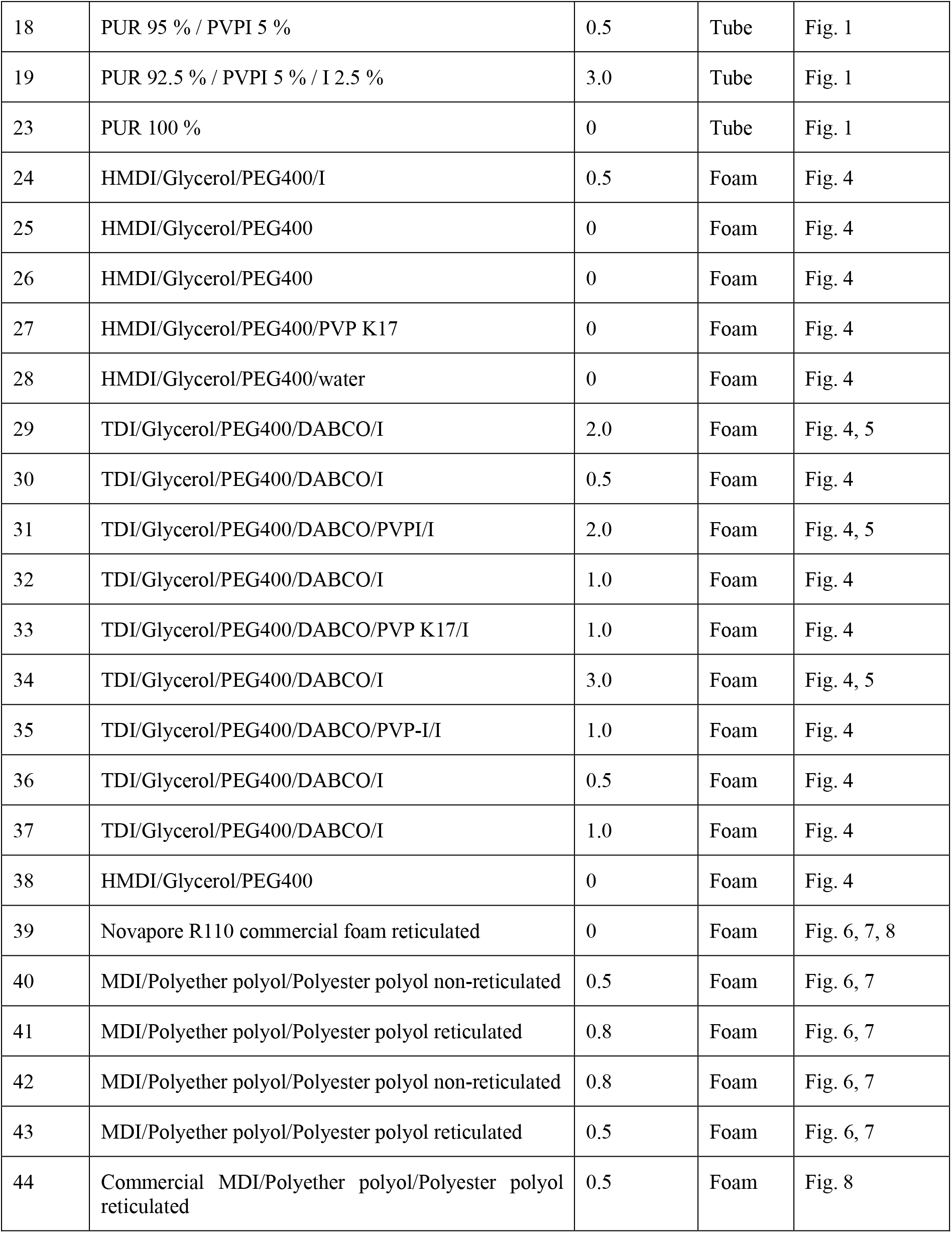
Composition and numeration of the tubes and foams used in this study.

**Figure 1.**
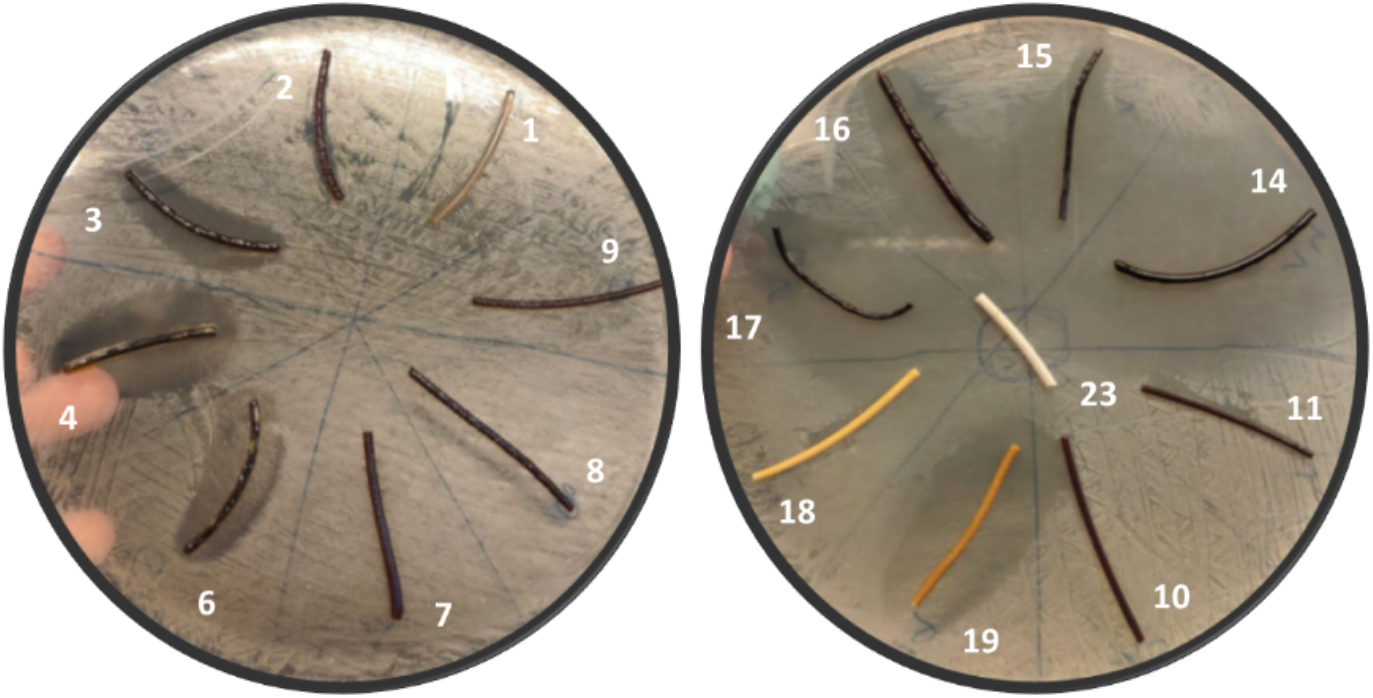
Inhibitory effects of PA- and PU-I thermoplastic tubes on growth of *S. aureus*. Materials 3, 4, 6, 14, 15, 16, 17 and 19 show clear growth inhibition zones when placed on a plate with *S. aureus* SH1000 (MSSA).

Based on the test results in **Figure 1**, two main observations can me made: (1) the size of inhibition zones is related to the total iodine loading. Generally, higher iodine loading will lead to increased activity. However, by observing and comparing tubes 6, 7, 8 and 9 that all possess an iodine loading of 5% a 2^nd^ observation can be made, (2) the compositional makeup of the PA-I complex is also important to determining biocidal activity.

Tubes 1, 2, 3, 4, 6 and 14 were selected to further evaluate the inhibitory effects on *S. aureus* USA300 (MRSA), *Streptococcus pyogenes, Staphylococcus epidermidis* ATCC 3598, *Candida albicans* ATCC 10231 and *Aspergillus fumigatus* 293 (**Figure 2**). A clear growth inhibition of all investigated microorganisms was observed with tubes 4 and 14 containing 10% iodine loading. Though the P-I complexes show activity against all indicated microorganisms, the biocidal effect is not always to the same level. Defined clearing zones were observed for tubes 3 and 6 at 5% iodine loading for *S. aureus USA 300*, but not for *S. pyogenes*. It should be noted that the plating studies utilized a high concentration of microorganisms. Such contamination concentrations would never likely occur in an actual healthcare setting. Expected contamination would occur at significantly lower concentrations and specific to the medical device surface, where significantly lower levels of iodine loading will probably suffice to reduce the infection risk.

**Figure 2.**
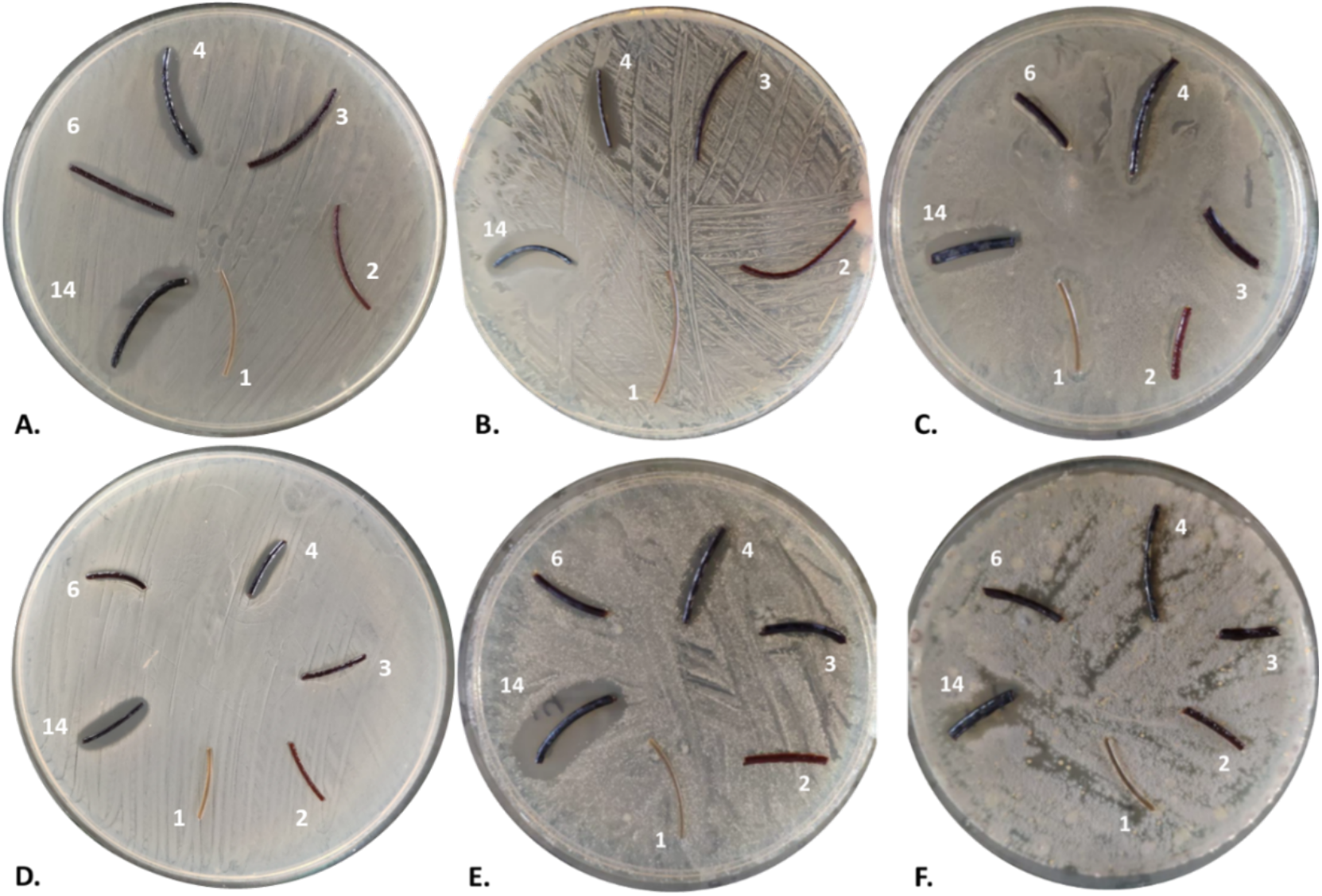
Inhibitory effects of PA-I tubes on various microorganisms. **A**. *S. aureus* SH1000 MSSA. **B**. *S. aureus* USA 300 MRSA. **C**. *S. pyogenes* clinical isolate. **D**. *S. epidermidis* ATCC 35984. For both *S. aureus* strains (A, B), *S. pyogenes* (C) and *S. epidermidis* (D) clear inhibition halos were observed with materials 4 and 14 as compared to the iodine-free control material 1. **E**. *C. albicans* ATCC 10231. Clear inhibition halos were observed for materials 4, 14 and a less clear inhibition halo for material 6. **F**. *A. fumigatus* 293, a clear inhibition halo was observed for material 14, and a smaller inhibition halo for material 4.

### 2.3. Inhibitory effects of PA-I tubes following sterilization

To evaluate the possibilities for re-use of the different materials, a potential decrease of the inhibitory effect following sterilization was tested. For both strains of *S. aureus* (MRSA and MSSA), the formation of halos reflecting growth inhibition was still observed upon autoclaving (121 °C, 15 psi, 15 min) (**Figure 3**). This observation also strongly supports the view that antibacterial activity for the iodine containing materials will be observed over an extended time period, which is an important factor for medical devices that need to remain in contact with the body for extended periods of time (e.g. wound dressings, urinary catheters, drains, etc.).

**Figure 3.**
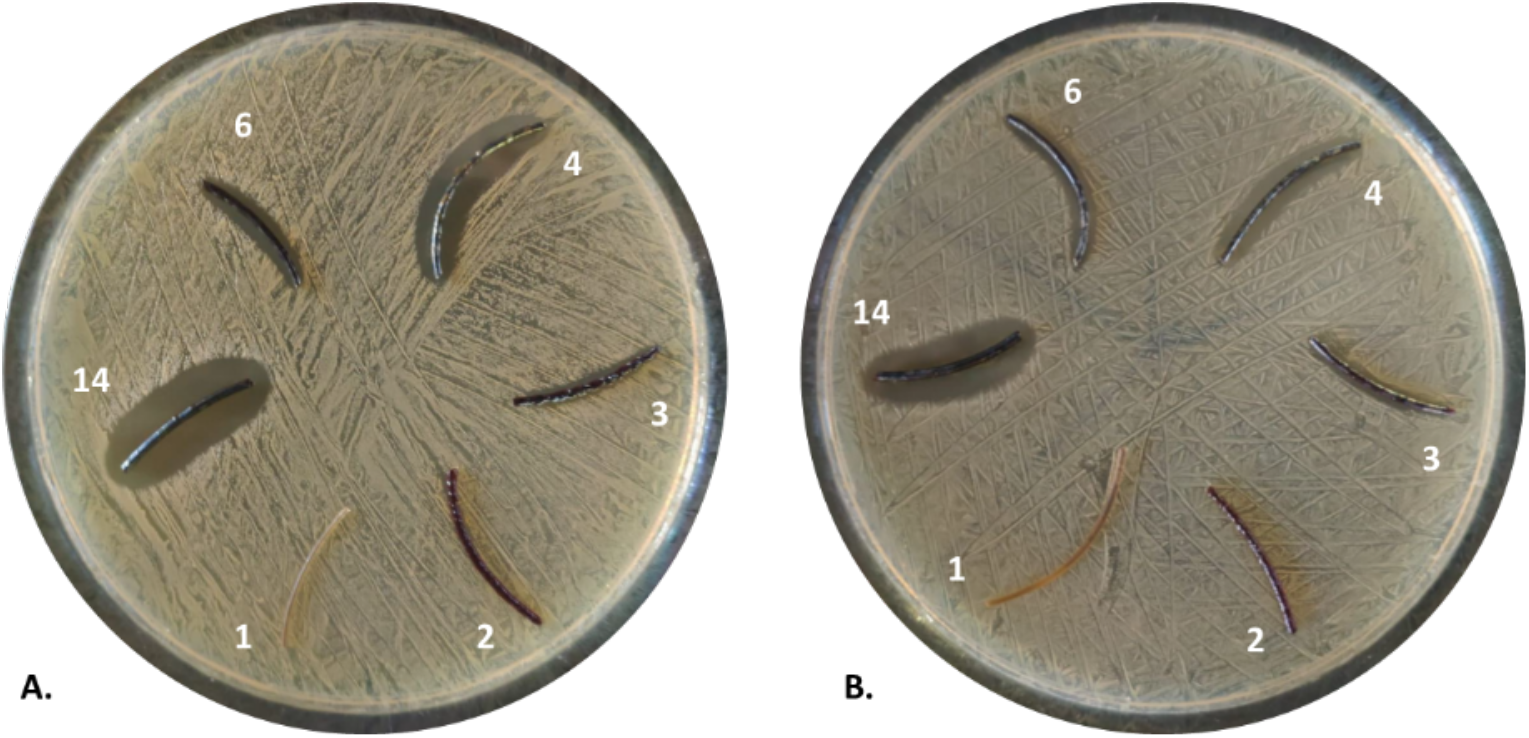
Growth inhibitory effects of PA-I tubes upon sterilization. **A**. *S. aureus* SH1000 MSSA. **B**. *S. aureus* USA 300 MRSA. For both *S. aureus* strains a clear inhibition halo was observed for the sterilized materials 4, 6 and 14 as compared to the sterilized iodine-free control material 1.

### 2.4. Inhibitory effects of PU-I foams on the growth of various microorganisms

To validate the antimicrobial effects of PVP-I and/or iodine in a different polymer context, PU-I foams with various loadings of iodine were included in the study. The PU-I foams were prepared as outlined in the materials and methods section. The inhibitory effects on microbial growth of lab-produced foams composed of different formulations (**Table 1**) were first tested on a clinical *S. aureus* isolate and the *S. epidermidis* type strain ATCC 3598. Foams 29, 31 and 34 showed clear inhibition halos (**Figure 4**) and they were, therefore, further evaluated for inhibition of *C. albicans* ATCC 10231 and a clinical strain of *S. pyogenes* that had been obtained after sonication of an implanted catheter (**Figure 5**).

**Figure 4.**
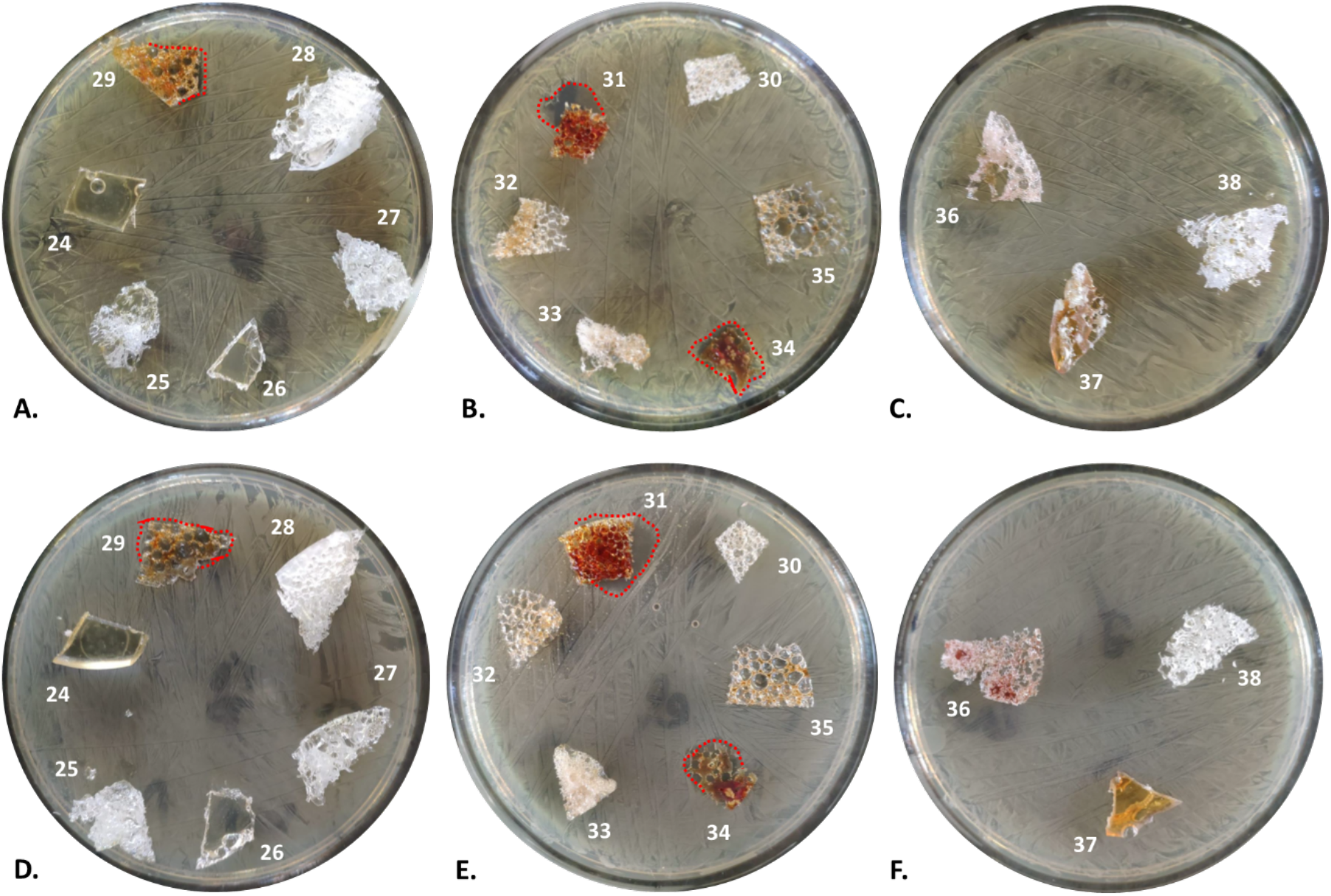
Growth inhibitory effects of laboratory PU-I foams on bacterial growth. **A, B** and **C**. *S. aureus* clinical isolate. **D, E** and **F**. *S. epidermidis* ATCC 35984. For both *S. aureus* and *S. epidermidis*, a clear inhibition halo was observed with materials 31 and 34. Material 29 caused a small growth inhibition halo. Inhibition halo’s are marked with a red dotted line.

**Figure 5.**
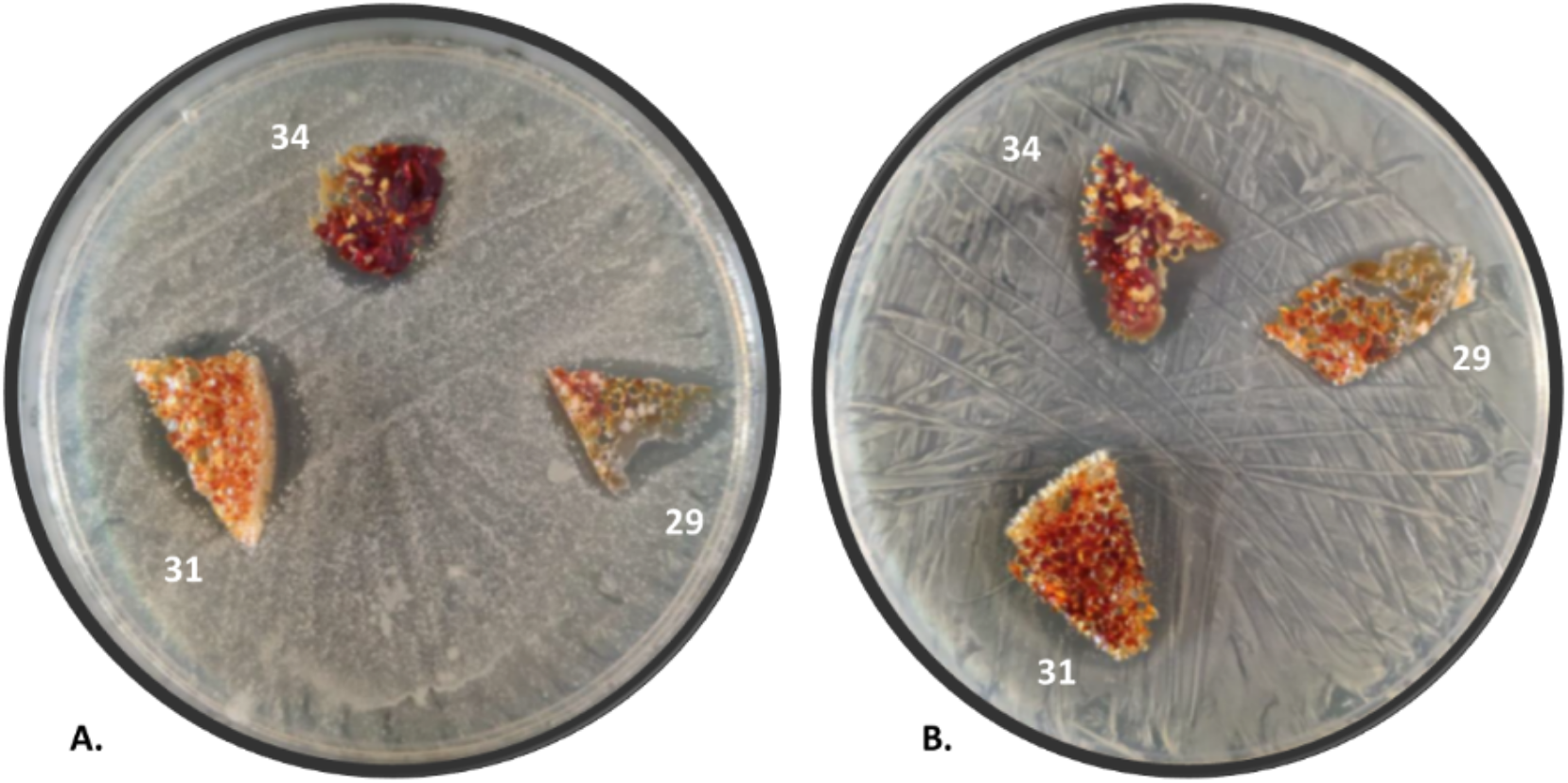
Growth inhibitory effects of selected laboratory PU-I foams on yeast and *S. pyogenes*. **A**. *C. albicans* ATCC 10231. **B**. *S. pyogenes* clinical isolate. For both microorganisms, clear inhibition halos were observed with all tested materials (29, 31, 34).

Based on the positive results from various lab-produced PU-I foams, new development foams, both reticulated and non-reticulated, were produced at pilot relevant scale (39-43, Table 1) and further evaluated against *A. fumigatus* (**Figure 6**). Due to the lower iodine loading in these foams, the fungal inhibition halo under the foams was evaluated to determine activity. Noteworthy is the observation that the fungicidal activity reflected by the clearing zones seems less related to the iodine levels, but more to the actual contact of the foams with *A. fumigatus* on the growth medium. In this respect, it is relevant that the light and buoyant PU-I foams do not all have the same contact points, and that the foams are either reticulated (open cell) or non-reticulated (closed cell). Accordingly, capillary effects of liquid can have differential effects of the observed inhibitory zones. This observation implies that the compositional makeup of the PU foam with respect to wetting, can also have indirect effects on biocidal activity. This parameter should be considered in medical device design.

**Figure 6.**
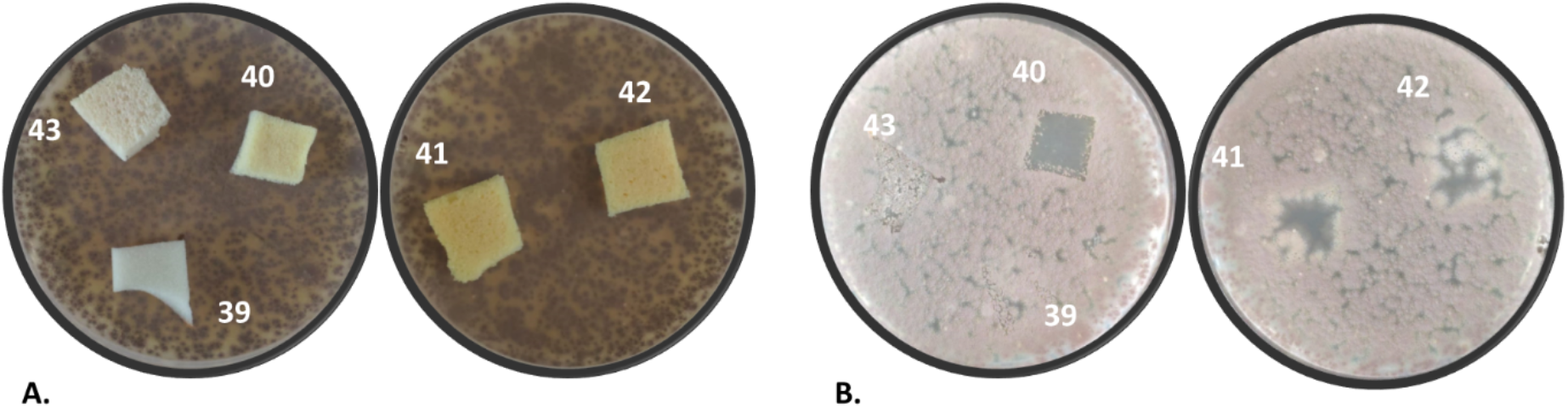
Growth inhibitory effects of development foams on *A. fumigatus*. **A**. Second generation foams 39 to 43 on top of MHA plates. **B**. Respective inhibition halos from foams 39 to 43. Clear halos were observed with foams 40, 41 and 42 when compared to the iodine-free control 39.

### 2.5. Inhibitory effect of PU-I foams on saliva samples from healthy volunteers

Due to the current great need of face masks, the developed foams were further evaluated for inhibition of oral microorganism. Freshly prepared and identical autoclaved foams 39 to 43 (**Table 1**) were incubated with saliva from healthy volunteers for 1 or 7 days at room temperature. Subsequently, the foams were replica-plated by pressing onto blood agar plates. Foams 40 to 43 showed a clear decrease in the microbial load compared to the control foam (39) after 1-day incubation of the saliva-inoculated material (**Figure 7A-B**). After 7 days incubation, the inhibitory effect was nearly 100% for the non-autoclaved foams (**Figure 7C-D**). Upon autoclaving, the foams retained their biocidal activity to varying extents.

**Figure 7.**
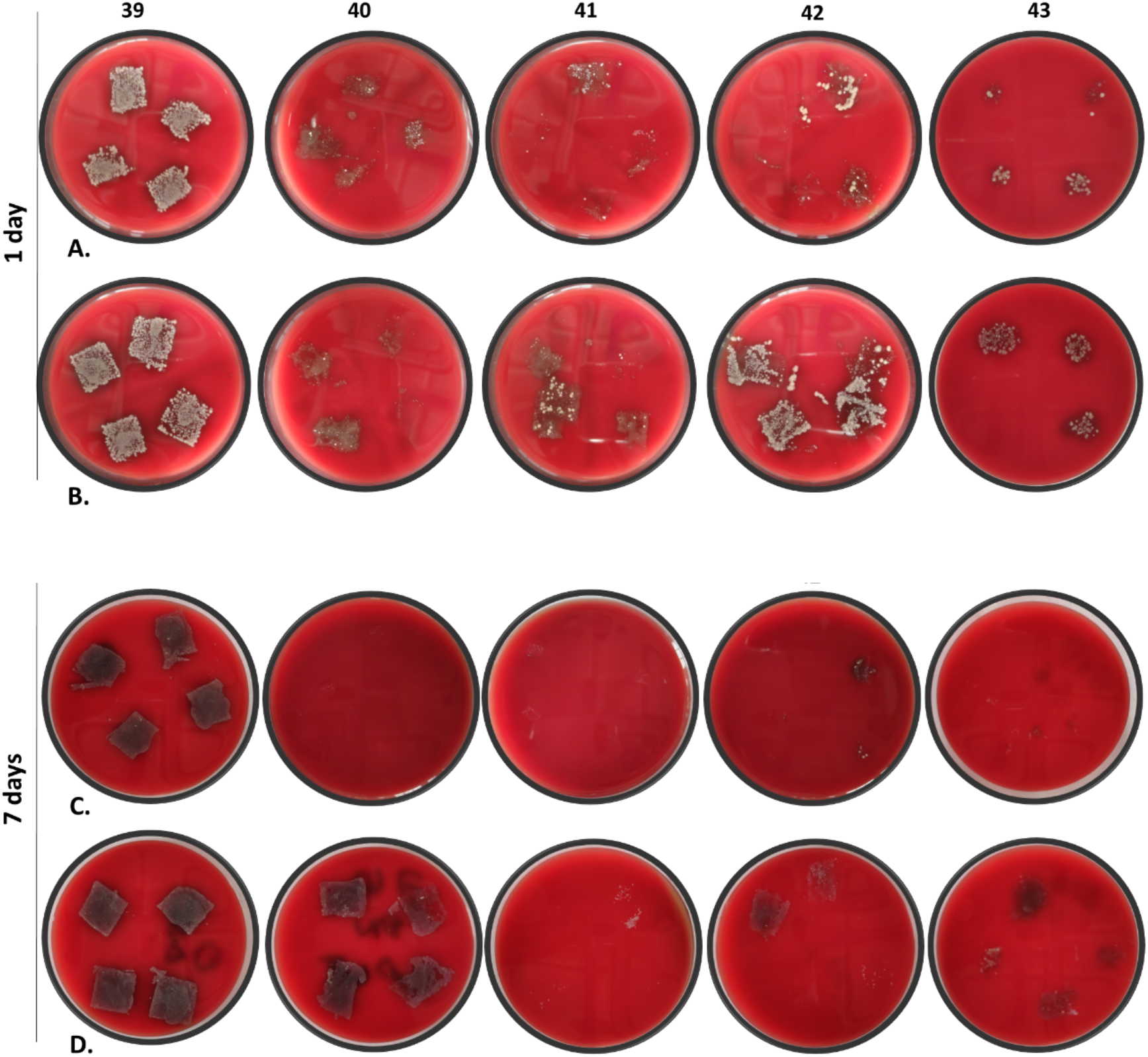
Microbial growth inhibition upon pre-incubation of various foams with saliva for 1 or 7 days. **A. and C**. Microbial load visualized by pressing foams 39 to 43 for 10 s onto a blood agar plate and subsequent overnight incubation. All foams show a clear inhibition of microbial growth compared to the iodine-free control 39. **B. and D**. Foams 39 to 43 were autoclaved prior the experiment and the microbial load was examined as in A and C.

### 2.6. Inhibitory effect of commercially produced PU-I face masks on microbial growth

Based on the above positive results, a commercial batch of reticulated foam 44 was produced at a batch size of 1500 kg in order to show the technology was scalable and commercially viable. The resultant high-quality foam was tested as a potential component for the development of face masks with microbial inhibitory properties. These face masks were worn for 3 h by healthy volunteers. Subsequently, the side of the foam that was in contact with the volunteer’s face was pressed on a blood agar plate to assess the bacterial load. As shown in **Figure 8**, the microbial load of face masks with foam 44 was significantly decreased in comparison with the control 39 (reticulated commercial foam with no biocidal activity).

**Figure 8.**
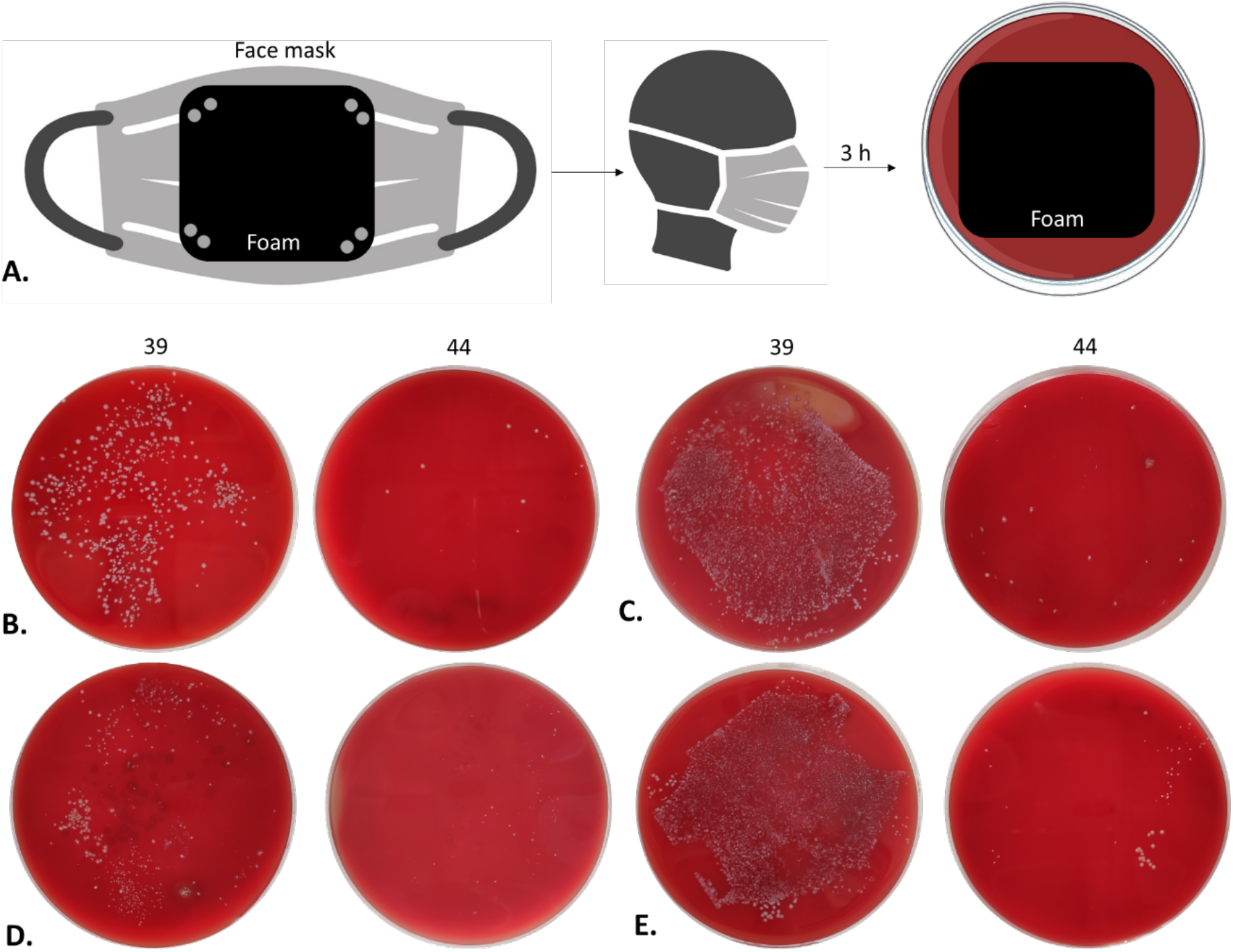
Microbial growth-inhibiting effects of PU-I foam face masks. **A**. Schematic representation of methodology (created in BioRender.com). **B**. Face masks made with foams 39 (control) or 44 (0.5% iodine) were worn for 3 h by a healthy volunteer and replica plated for 10 s on top of a BA plate, or **C**. kept on top of a BA plate during 24-h incubation at 37 °C. **D**., **E**. Respective A and B results after washing the face mask 1x at 60 °C prior usage.

## 3. Discussion

Considering the impact of HAI’s in many clinical settings, developing medical devices with inherent properties against bacterial contamination and biofilm formation is of great importance. Many studies are focused on the evaluation of coating agents, such as polymer coatings, antimicrobial coatings or nanostructured coatings, to antagonize the first adherence of microorganisms and, thus, inhibit or slow down biofilm formation ^[13]^. From such studies, the use of antiseptic-based coatings was shown to be more effective when compared to the use of antibiotics-based coatings ^[13, 14]^. Moreover, the local delivery of antiseptics was successful when implemented in coatings for catheter-based applications ^[15, 16]^. However, antiseptics often pose an unknown risk once placed in the body due to their synthetic and toxic nature.

Iodine is a naturally occurring element shown to have broad antimicrobial and antiviral activity, but it is also an essential mineral nutrient. Iodide salts are often added to foods and supplements to ensure good human health. Elemental iodine, itself, shows poor water solubility, is corrosive and readily sublimes (unstable physical form), which has limited its use as a biocide until the discovery of the povidone-iodine complex, the active material in Betadine^®^. The povidone-iodine complex is: stable, water-soluble, highly active against a broad range of microorganisms and viruses, and safe for human use. As such, the povidone-iodine material has become an essential tool for the prevention of infections with broad use in both institutional and public healthcare. Though povidone-iodine represents a true success story for harnessing the antimicrobial and antiviral properties of iodine for water-based applications, there has been relatively limited success in harnessing said iodine attributes in water-insoluble engineering plastics that could be used for the fabrication of various medical devices, and healthcare and consumer products.

In the present study, we show that the excellent antiseptic properties of water-soluble povidone-iodine, can be similarly realized in water-insoluble, engineering plastics, specifically PA- and PU-iodine complexes. The PA- and PU-iodine complexes: can be manufactured cost effectively, have desirable mechanical properties and are inherently antibacterial. Such materials are expected to have broad-use applications in the fabrication of antibacterial/antiviral articles for application in healthcare, consumer care and industry.

The PA- and PU-iodine complexes show activity against a broad range of pathogens, which will have a direct effect of reducing infections by inhibiting microbial growth due to contamination or reducing undesirable biofilm formation on products made from said materials. The complexation of iodine to polymers that are commonly used to fabricate a range of medical devices results in materials with reduced infection risk, while maintaining the safety-profile similar to povidone-iodine. Our results provide proof-of-principle that PA- and PU-I complexes can be produced that: are commercially viable, show broad biocidal activity and have broad-use application. The antiseptic attributes of PVP-I were successfully integrated in two commonly used biomaterial families, polyamides and polyurethanes, by the complexation with iodine. We also show that the level of iodine concentration or the incorporation of other ingredients in the systems can have a significant effect on the growth inhibitory zones, suggesting that the polymer-iodine systems can be further customized to meet specific requirements for a particular medical need. Moreover, we show that the biocidal properties of the polymer-iodine complexes included in this study are largely maintained upon sterilization, which strongly supports that the materials can be re-used or will maintain their biocidal activity for an extended period of time. Thus, these polymers have the extra advantage of a low environmental footprint, since they can be recycled or allowed to remain in use for longer periods of time.

In the case of PU-I foams, we show that the technology is readily scalable and that commercial production of both non-reticulated and reticulated foams with biocidal activity is viable. In order for the PU-I foams to have broad-use applications, it is important that both non-reticulated and reticulated foams can be produced. In non-reticulated PU foams, the gas bubbles entrapped in the foam are completely closed and thus air or liquid cannot pass through the foam structure. Such non-reticulated foams are generally used for their cushioning/insulation properties, but cannot be used in filters. To make the PU foam acceptable for filter applications, the foam must be reticulated, which is the process of treating the corresponding non-reticulated foam with a combustible gas or chemical to bring about the controlled degradation of the gas bubbles to produce an open and highly porous foam structure. Once the PU foam is reticulated, the foam can be used for the production of efficient filters. The present research shows that the biocidal activity observed in the non-reticulated PU-I foam is maintained after reticulation. Finally, we developed a prototype face mask with a commercially produced PU-I reticulated foam that shows strong biocidal activity against oral bacterial flora. Such reticulated foams can be implemented in both face masks and filters of air handling systems to reduce the spread of airborne pathogens.

## 4. Conclusion

In conclusion, the beneficial antimicrobial attributes of the water soluble PVP-I can be similarly realized in PA- and PU-I materials. The production process of such P-I plastics requires no additional complex processing and existing manufacturing infrastructure can be utilized, which makes them commercially viable. The P-I plastics can be fabricated to generate a host of medical devices that are inherently less susceptible to infection generation and commonly used in healthcare practice. The production of engineering plastics that release iodine in a controlled and safe manner, allows for the fabrication of existing medical devices with reduced infection risk and opens new treatment options for a wide range of applications. Both PA and PU materials show excellent biocompatibility and thus are used for the fabrication of a broad range of medical devices. By introducing biocidal properties in a safe manner through the complexation of iodine, one can continue to fabricate urgently needed medical devices that reduce the risks of surgical site- and other hospital-acquired infections. Importantly, the implications of this technology could be more far-reaching than just healthcare-related applications, since the base materials are also used in many consumer and industrial applications where antimicrobial properties would be a favorable attribute to decrease the spread of infections, such as in carpets, air filters and protective face masks.

## 5. Experimental Section/Methods

### Preparation of the tubes and foams

Thermoplastic PA and PU systems (samples 1-23) were prepared via hot melt extrusion, using a Thermo Prism Eurolab 16 twin screw extruder having 16 mm screw diameter and 25 cm barrel length. The extruding barrel had five heating zones that were set in the range of 150-240 °C depending on the system being compounded. The rotation speed of the screws was fixed at 400 rpm. The powders/solids were initially dry mixed and then fed into the extruder. The extruded filament was water-cooled and cut to length to give the tubes. The following abbreviations are used:

- PA6: Polyamide 6, Lanxess Durethan^®^ 29
- PA6 econyl: Polyamide 6, Aquafil ECONYL^®^ 6
- PVP: Polyvinylpyrrolidone K30, Boai NKY Pharmaceuticals Ltd. PolyViscol™ K30
- PVPI: Polyvinylpyrrolidone-iodine, Boai NKY Pharmaceuticals Ltd. KoVidone^®^-I
- PA11: Polyamide 11, Sigma Aldrich/Merck
- PA12: Polyamide 12, Sigma Aldrich/Merck
- PUR: aliphatic polyether thermoplastic PU, Lubrizol TecoFlex™ EG-93A-B30
- I: elemental iodine 99.8%, Sigma Aldrich/Merck

Thermoset PU systems (samples 24-38) were prepared by the reaction of an isocyanate with polyethylene glycol/glycerol polyol blend with or without an iodine source. If an iodine source was used in the reaction, the iodine was initially dissolved in the polyol blend prior to conducting the polyurethane reaction. Polyurethane reactions including an iodine source required the addition of a catalyst (DABCO) to accelerate the reaction. The polyurethane reaction was conducted by reacting the isocyanate with the polyol source at room temperature until homogeneity, and the reaction mixture was then poured into molds and placed in a 60 °C oven for 2 hours. The following abbreviations are used:

- HMDI: Hexamethylene diisocyanate, Sigma Aldrich/Merck
- TDI: Toluene diisocyanate, Sigma Aldrich/Merck
- PEG400: Polyethylene glycol 400, Sigma Aldrich/Merck
- Glycerol: Sigma Aldrich/Merck
- PVP K17: Polyvinylpyrrolidone K17
- DABCO: 1,4-diazabicyclo[2.2.2]octane Sigma Aldrich/Merck

Thermoset PU systems (samples 39-44) were kindly prepared by Caligen Europe BV. The PU systems were based on commercial processes and raw materials, and supplied in their corresponding non-reticulated and reticulated forms. The reticulation process was conducted by combustion. Sample 39 is a commercial reticulated foam containing no iodine (i.e. control). Samples 40-43 were lab samples based on MDI (methylene diphenyl diisocyanate)/polyether polyol/polyester polyol/iodine foams that were produced on the kg scale. Sample 44 is a reticulated foam based on MDI/polyether polyol/polyester polyol/iodine commercially produced at a batch size of 1500 kg.

### Inhibition effect of PA/PU-I tubes/foams on microbial growth

Methicillin-susceptible *Staphylococcus aureus* (MSSA) SH1000, methicillin resistant *S. aureus* USA 300 (MRSA), a clinical *Streptococcus pyogenes* isolate, *S. epidermidis* ATCC 35984, *Candida albicans* ATCC 10231, and *Aspergillus fumigatus* 293 were grown in tryptic soy broth (TSB) overnight at 37 °C. 100 µl of the overnight cultures were diluted to an optical density at 600 nm (OD_600_) of 0.1 and subsequently plated on 100 mm Mueller-Hinton agar plates (MHA). Tubes or foams with different concentrations of nylon, PVP and iodine were placed on top of the plates, and the plates were then incubated at 37 °C. Growth inhibition was inspected upon 24 and 48 h of incubation.

### Microbial growth inhibition by PA-I tubes upon sterilization

To evaluate a possible decrease in the microbial growth inhibitory efficacy of the investigated materials upon standard sterilization by autoclaving, the tubes were washed once with 70 % ethanol, rinsed with distilled H_2_O, dried and autoclaved at 121 °C (15 min, 15 psi).

### Growth-inhibiting effects of PU-I foams on microbe-containing saliva samples from healthy volunteers

To evaluate the effects of PVP-I on oral microorganisms, 1 mL saliva samples from healthy volunteers were incubated with various foams (materials 39 to 43, Table 1). After 1 or 7 days of incubation at room temperature, the foams were replica plated by gently pressing them onto Blood Agar plates (BA) for 10 s. Subsequently, the plates were incubated for 24 h at 37 °C. To ensure that all the microorganisms present in the foams were detected, the materials incubated with saliva for 7 days were also plated and kept on top of the BA plates during incubation for 24 h at 37 °C. Evaluation of the growth inhibitory effects of 1x autoclaved foams was carried out in parallel.

### Microbial growth-inhibiting effects of PU-I foam mouth masks

Prototype face masks were developed with commercial PU foams containing iodine (sample 44) and without iodine (sample 39) to evaluate the activity of these foam materials on microorganisms commonly transmitted from the face and breath to face masks. Mouth masks were worn for 3 h by volunteers. Afterwards, the inside of the mask was gently pressed for 10 s on BA plates. The plates were subsequently incubated for 24 h at 37 °C. After pressing the inside and outside of the mask on BA plates, a piece of the mask was cut and incubated on top of a BA plate at 37 °C for 24 h.

### Ethical approval

Studies with healthy volunteers wearing PU-I foam face masks were performed based on written informed consent with approval of the Medical Ethics Review Board of the University Medical Center Groningen (UMCG; approval no. Metc2012-375), and with adherence to the Helsinki Guidelines.

## Acknowledgements

We would like to acknowledge the help and assistance of Caligen B.V./Vita Group for the manufacture and supply of the development and commercial PU-I foams tested in this study.

## Author contributions

ML-A, HU, NK and JMvD: conceptualization; ML-A and HU: data curation; ML-A and HU: formal analysis; HU, NK and JMvD: funding acquisition; ML-A and HU: investigation; ML-A, HU and JMvD: methodology; HU, NK and JMvD: project administration; HU, NK and JMvD: resources; HU and JMvD: supervision; ML-A and HU: validation; ML-A: visualization; ML-A: writing - original draft; ML-A, HU, NK and JMvD: writing - review & editing. Data and materials availability: All data and materials are available.

